# Subcellular Organization of Viral Particles During Maturation of Nucleus-Forming Jumbo Phage

**DOI:** 10.1101/2021.04.26.441357

**Authors:** Vorrapon Chaikeeratisak, Kanika Khanna, Katrina T. Nguyen, MacKennon E. Egan, Eray Enustun, Emily Armbruster, Kit Pogliano, Elizabeth Villa, Joe Pogliano

**Affiliations:** Division of Biological Sciences, University of California, San Diego, CA 92093, USA; Department of Biochemistry, Faculty of Science, Chulalongkorn University, Bangkok, 10330, Thailand

## Abstract

Many eukaryotic viruses assemble mature particles within distinct subcellular compartments, but bacteriophages were long assumed to assemble randomly throughout the host cell cytoplasm. Here we visualized the subcellular location of viral particles formed during replication of *Pseudomonas* nucleus-forming jumbo phages and discovered that they assemble a unique structure inside cells we term phage bouquets. We show that after capsids complete DNA packaging at the surface of the phage nucleus, tails assemble and attach to the capsids, and these particles accumulate to form bouquets at specific subcellular locations. In these bouquets, the viral particles are arranged in a spherical pattern with tails oriented inward and the heads outwards. Localized at fixed distances on either side of the phage nucleus, bouquets grow in size and number over time as new phage particles are added. In the presence of mutations that cause the phage nucleus to be mispositioned away from its typical position at the midcell, bouquets still localize at the same fixed distance from the nucleus, suggesting an active mechanism for their formation and positioning. These results mark the discovery of a pathway for organizing mature viral particles inside bacteria and demonstrate that nucleus-forming jumbo phage, like most eukaryotic viruses, are highly spatially organized during all stages of their lytic cycle.

## Introduction

Subcellular organization is an important trait in all domains of cellular life, allowing cells to carry out their myriad required functions faithfully and efficiently, and also plays a key role in the replication of viruses [1-4]. Upon infection of a host cell, many eukaryotic viruses manipulate host proteins and components of various organelles such as the Golgi apparatus and endoplasmic reticulum to form replication factories in which they can safely and efficiently replicate their genomes [5-9]. They also assemble mature viral particles in specific subcellular compartments [1, 10]. Despite bacteria generally lacking organelles to manipulate and the longstanding assumption that bacteriophage replication is spatially disorganized, recent work suggests that some phages reproduce by forming replication factories and display a high level of subcellular organization within their host cells [11-20].

We recently demonstrated that Pseudomonas jumbo phage replication is spatiotemporally organized and involves formation of a key structure called the phage nucleus [11, 12, 21]. Immediately upon injection of phage DNA into the host cell, several large bacteriophages including related *PhiKZ-like Pseudomonas* phage PhiPA3, 201Phi2-1, and PhiKZ, as well as *Serratia* phage PCH45, enclose their replicating DNA in a roughly spherical, nucleus-like, proteinaceous shell that expands as the viral DNA replicates [11-13, 21-23]. The phage nucleus compartmentalizes proteins according to function, with proteins involved in DNA replication, recombination and transcription localized inside and those involved in translation and metabolic processes outside [11, 12, 24]. This compartmentalization provides an advantage to nucleus-forming phages by allowing them to shield their DNA from host-encoded DNA restriction and CRISPR/CAS systems [13, 25]. The bacterial chromosome is degraded by an as yet to be identified nuclease, leaving the bacterial cell unable to further mount a genetic response to the phage infection [11, 12, 22, 23]. Early during infection, filaments of the tubulin-like phage protein PhuZ assemble at each cell pole [22, 23]. Over time, these filaments extend and push the phage nucleus along the cell length to ultimately form a bipolar spindle that positions the phage nucleus at the midcell [22, 23]. The PhuZ spindle’s dynamic instability allows the filaments to grow and shrink in order to keep the phage nucleus centered at midcell throughout infection [22, 23]. Later during the infection cycle, capsids assemble on the cell membrane and migrate along the treadmilling PhuZ filaments to reach the surface of the phage nucleus, where they dock and initiate DNA packaging [21]. As it traffics the capsids to the phage nucleus, the PhuZ spindle also pushes on the phage nucleus, causing it to rotate at midcell [11, 21]. Notably, elements of this sophisticated spatiotemporal subcellular organization, including compartmentalization of viral replication and transport of viral components along cytoskeletal elements, mirror those of many eukaryotic viruses [26-28].

Spatial organization and macromolecular assembly can also play important roles in viral evolution. By studying phages PhiPA3 and PhiKZ, we identified Subcellular Genetic Isolation and Virogenesis Incompatibility as two general mechanisms that contribute to viral speciation [29]. Subcellular Genetic Isolation reduces the ability of two viruses that have entered the same cell to undergo genetic exchange because they are spatially segregated [29]. For nucleus-forming jumbo phages, co-infections can result in Subcellular Genetic Isolation due to the establishment of two separate nuclei that physically isolate the two competing viral genomes [29]. Virogenesis Incompatibility occurs when common components from two different phage interact negatively to inhibit viral particle production during co-infection of the same host cell [29]. In the case of PhiPA3 and PhiKZ, for example, PhuZ monomers from each phage are similar enough to each other to co-assemble but due to their divergence, form nonfunctional hybrid polymers that disrupt phage nucleus positioning and capsid trafficking to the phage nucleus and therefore reduce phage DNA packaging. Subcellular Genetic Isolation and Virogenesis Incompatibility are likely widespread speciation mechanisms among eukaryotic and prokaryotic viruses and highlight the importance of understanding the mechanisms underlying viral spatiotemporal subcellular organization.

While we are now beginning to appreciate the subcellular complexity of nucleus-forming bacteriophages and its potential effects on viral speciation, less is known about the final stages of phage particle assembly. The assembly and maturation of many eukaryotic viruses are known to occur in defined subcellular compartments [1-9]. In contrast, the major steps of tailed phage maturation, which involves separate assembly of the phage tail and capsid, followed by viral DNA packaging and finally attachment of the phage head and tail, has been observed proceeding in an apparently random manner throughout the cell, leading to a random distribution of complete phage particles filling the host cytoplasm before lysis [30-33]. Although there have been reports of phage capsids clustering in the host cytoplasm, the cause of this phenomenon and its possible role in phage assembly have not been investigated [34-36].

As we have observed at all other stages of their replication cycle, we show here that large, nucleus-forming *PhiKZ-like* viruses achieve a high level of subcellular organization during the final stages of phage maturation. Using fluorescence microscopy and cryo-electron tomography (cryo-ET), we demonstrate that apparently complete viral particles are not randomly distributed throughout the cell but instead form micelle-like spheres with phage tails in the center and heads pointing outwards, reminiscent of a bouquet of flowers. This marks the first discovery of highly organized bacteriophage particle assembly *in vivo* and demonstrates that nucleus-forming jumbo phages are spatially organized during all stages of the lytic cycle, including the final stages of phage assembly. These findings, together with other recent studies [11, 12, 14-23, 29], establish a new paradigm of phage subcellular organization that requires further study to understand the underlying mechanisms and possible evolutionary benefits. It also demonstrates that subcellular organization of virion assembly is conserved among viruses infecting different kingdoms of life.

## Results and Discussion

We investigated the major steps of viral maturation in nucleus-forming jumbo phage PhiPA3 using fluorescence microscopy. *P. aeruginosa* cells were infected with PhiPA3, stained with DAPI to reveal DNA, and imaged throughout the infection cycle. Using fluorescence microscopy and FISH, we have previously demonstrated that *Pseudomonas* phage PhiPA3, 201Phi2-1 and PhiKZ degrade the host chromosome early in the infection cycle, which allows the use of DAPI staining to visualize phage DNA. At 15 minutes post infection (mpi), during the early stages of PhiPA3 replication, DAPI staining revealed the phage nucleus as a small focus near one cell pole (Fig. 1 A, top leftmost panel, orange arrow), while the host chromosome filled most of the cell. By 30 and 45 mpi, the host chromosome had been degraded and the phage nucleus appeared as a larger focus located near midcell as previously reported (Fig. 1 A, top panels, white arrows) [23]. From 60 to 90 mpi, however, two new DAPI-staining structures appeared on each side of the phage nucleus. These structures appeared as foci at 60 mpi and as rings at 75 and 90 mpi (Fig. 1 A, top panels, green arrows). The rings only appeared after DNA packaging into capsid heads (60 mpi), suggesting that they are composed of capsids containing viral DNA assembled into a spherical structure that appears as a ring when viewed in cross section. The number of DAPI rings per cell increased over time, such that at 60 mpi, infected cells contained either one (3%, n= 126) or two rings (8%, n=126) of capsids (Fig. 1E). By 75 mpi, more than 50% of infected cells contained 2 rings (Fig. 1 A, top panels & Fig. 1E, n= 117), and by 90 mpi, more than 4 rings were detected in some cells (Fig. 1B & E, n= 128). We used structured illumination microscopy (SIM) (Fig. 1C) to observe these structures at higher resolution in DAPI stained infected cells 90 mpi. At this resolution, the rings appeared as two horseshoe-shaped structures that averaged ∼660 nm (n= 8) in diameter with an opening of ∼150 nm (n= 8) located ∼1 µm away from the phage nucleus when measuring from the center of the horseshoe structure to the center of the phage nucleus (Fig. 1C, n= 8, & 1F, n= 252). The distance from the edge of the horseshoe structure to the edge of the phage nucleus averaged ∼200 nm (n=255, Fig 1F).

**Figure 1.**
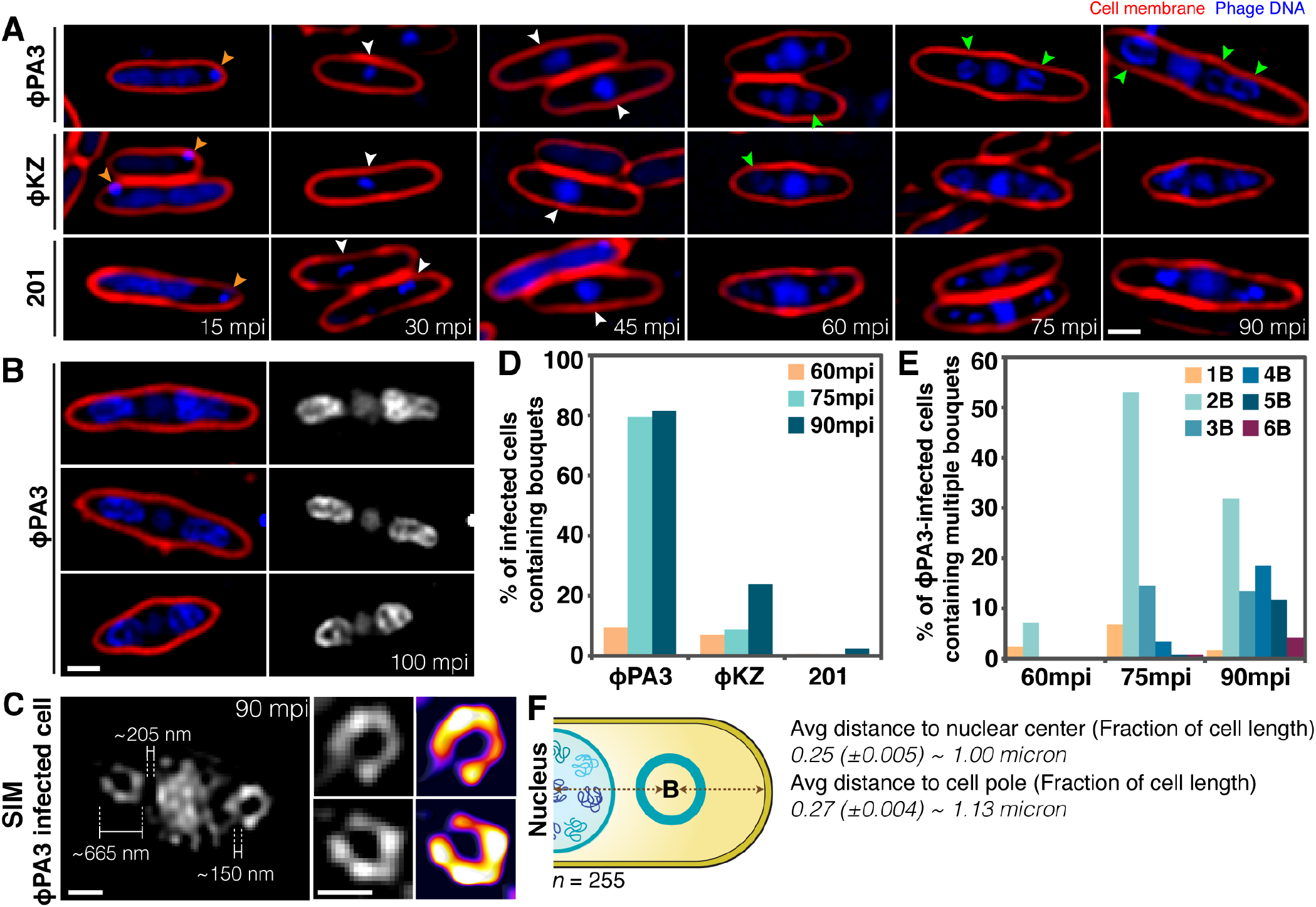
Maturation process and bouquet formation of phages PhiPA3, PhiKZ and 201Phi2-1. (A) Time-series images over a course of infection of *P. aeruginosa* by phage PhiPA3 (upper panels), PhiKZ (middle panels), and *P. chlororaphis* infected with 201Phi2-1 (bottom panels). Cell membranes were stained with FM4-64 (red) and DNA was stained with DAPI (blue or gray). The phage genome was injected into the host cell close to the pole (orange arrows) at early infection and bouquet structures appeared adjacent to the phage nucleus after 60 mpi (green arrows). (B) Still images of phage PhiPA3-infected *P. aeruginosa* cells at late infection. (C) 3D-SIM image of the bouquet structures in phage PhiPA3-infected *P. aeruginosa* cells at 90 mpi reveals horseshoe-shaped structures adjacent to the phage nucleus with average distance and size indicated (n of bouquets = 8). Subpanels show magnified images of the horseshoe-shaped structure and its heatmap intensity. (D) Graph showing the percentage of *Pseudomonas* cells infected with phages PhiPA3, PhiKZ and 201Phi2-1 containing bouquets at 60, 75, and 90 mpi. (E) Graph showing the percentage of phage PhiPA3-infected *P. aeruginosa* cells containing different number of bouquets **B** at 60, 75, and 90 mpi. (F) Average bouquet size and its distance from the nuclear center and cell pole (n = 255). The numbers were expressed as a fraction of cell area or cell length as indicated. Scale bars: 1µm (A,B,C insets), 0.5µm (C main panel).

These horseshoe shaped structures also appeared in host cells during infection by related nucleus-forming jumbo phage PhiKZ and 201phi2-1, albeit at a significantly reduced frequency. While over 80% of *P. aeruginosa* cells infected with phage PhiPA3 contained these structures at 75 mpi (Fig. 1D, n=117), they were observed less frequently in *P. aeruginosa* cells infected with phages PhiKZ (∼8% n=184) or *P. chlororaphis* infected with 201Phi2-1 (0.5%, n= 197) (Fig 1A, D). These results suggested that formation of these structures is conserved but likely not essential for nucleus-forming jumbo phage replication.

The extranuclear DAPI-stained structures suggested that DNA-filled capsids accumulated at specific subcellular locations late during infection. To test this hypothesis, we visualized localization of a GFP-fusion to capsid protein gp136 simultaneously with DAPI staining at various stages of PhiPA3 infection. At early time points, gp136-GFP appeared uniformly distributed in the cytoplasm, excluded from the phage nucleus (Fig. 2A, 30 mpi). At 45 mpi, gp136-GFP localized to the periphery of the phage nucleus, indicating capsids assembled and docked on the phage nucleus (Fig. 2A, 45 mpi). From 60 mpi to 100 mpi, capsids re-localized to distinct regions on each side of the nucleus, forming small foci that co-localized with the DAPI rings (Fig. 2A). Additionally, DAPI staining intensity in the flanking rings increased over time, while simultaneously decreasing in the phage nucleus (Fig. 2C). Together, these results support the conclusion that DAPI rings visualized at late stages of infection are composed of capsids that have been filled with DNA at the nucleus surface and then migrated to be arranged in spheres on each side of the nucleus.

**Figure 2.**
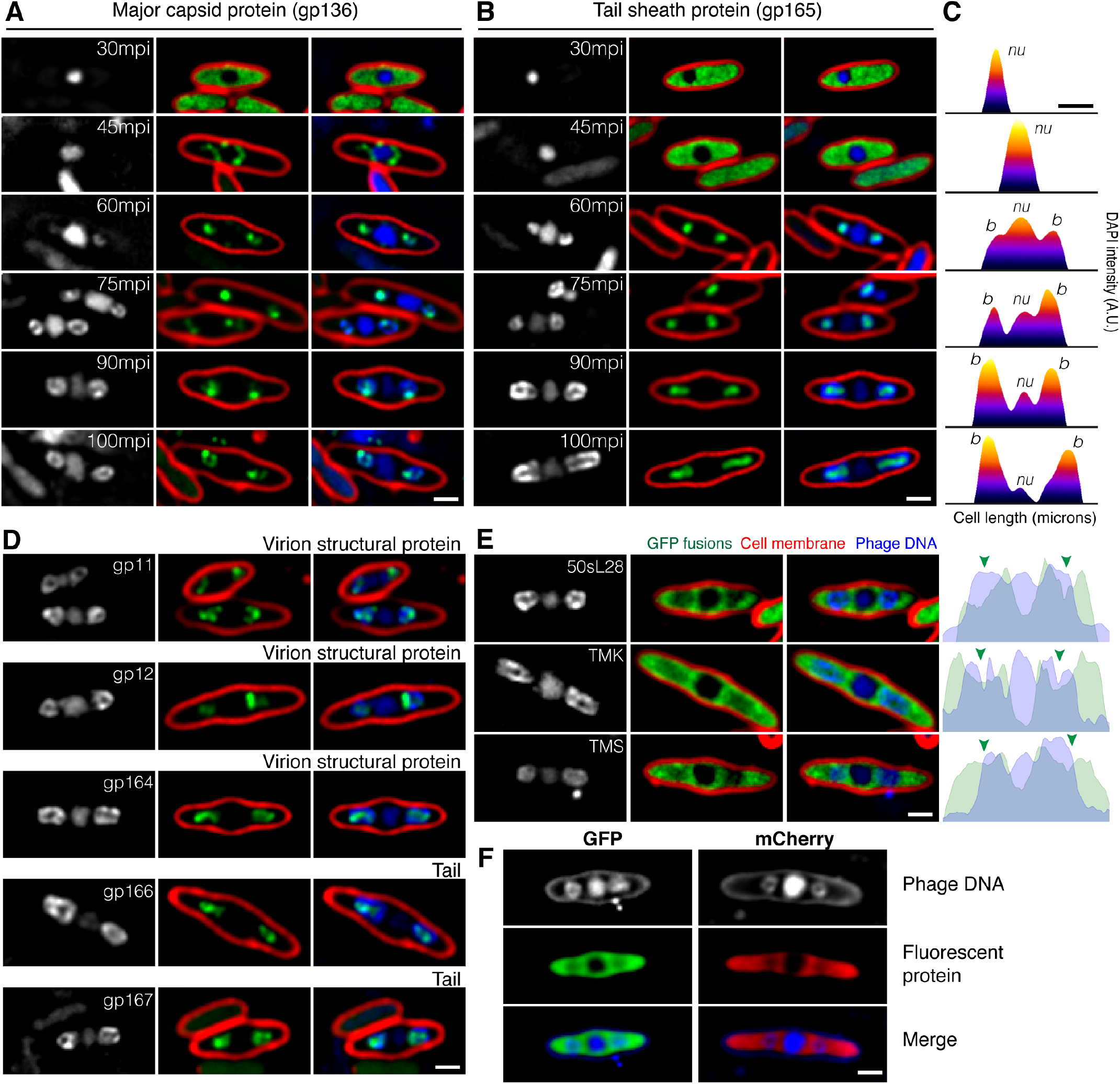
Localization of virion-structural proteins in phage PhiPA3-infected *P. aeruginosa* cells. The cells were grown on an agarose pad and expression of the GFP fusion protein (green) was induced by arabinose at indicated concentration; 0.01% for gp11 and gp12, 0.05% for gp136, gp165, 50sL28, TMK, and TMS, 0.10% for gp164, gp166, and gp167, 0.20% for GFP and mCherry. Phage PhiPA3 was added onto the pad to initiate infection. Cell membrane was stained by FM4-64 (red) and DNA was stained by DAPI (blue or gray). (A, B) Temporal localization pattern of major capsid protein gp136 (A) and tail sheath protein gp165 (B) during infection relative to the location of the phage nucleus and bouquets. (C) Heat maps of the DAPI intensity inside the PhiPA3-infected cells as corresponding to the fluorescent micrographs in (B) showing DNA distributed from the nucleus into bouquets at late infection. *nu* and *b* indicate the position of the phage nucleus and bouquets. (D) Localization of other structural proteins in PhiPA3-infected cells at 75 mpi: virion structural proteins gp11, gp12, gp164, and tail proteins gp166 and gp167. These proteins localized with DAPI indicating they are part of the capsid or accumulated inside the bouquets suggesting that they are tail proteins. (E) Localization of host ribosomal protein (50sL28) and metabolic enzymes (Thymidylate kinase, TMK, and Thymidylate synthase, TMS) in PhiPA3-infected cells at 75 mpi. The fluorescent intensity map (right panels) revealed the partial exclusion of the fusion protein (green) from the phage nucleus and bouquets (blue). Arrows indicate the location of bouquets. (F) Localization of soluble GFP and mCherry in PhiPA3-infected cells at 75 mpi. Scale bars: 1µm.

To determine the subcellular location of other virion structural components relative to the capsids, we GFP-tagged six additional PhiPA3 structural proteins, three of which are likely involved in tail assembly. The tail sheath protein (gp165) was uniformly distributed and excluded from the phage nucleus early during infection (Fig. 2B, 30 and 45 mpi) but at later time points, it localized inside the ring of capsids (Fig. 2B-C, 60-100mpi). At these late time points, the bright ring of DAPI staining capsids completely surrounded the fluorescence from GFP-tagged gp165. Three additional phage structural proteins (gp164, gp166, gp167) also localized inside the DNA rings with the other tail proteins (Fig, 2D), while gp11 and gp12 co-localized with the DAPI-stained capsid rings (Fig, 2D). For all GFP-tagged phage proteins, uninfected control cells showed that the proteins were uniformly diffused throughout the cytoplasm, with gp11 also forming small puncta (Fig. S1). These results suggest that, late during infection, viral particles assemble a spherical structure with heads located on the outside and the tails pointed inward. We term these structures phage “bouquets”, in reference to previous reports that describe clusters of phages observed when examining phage lysates by electron microscopy [37-39].

Using GFP fusions, we have previously shown that proteins such as thymidylate kinase (TMK-GFP), 50s ribosomal protein subunit (L28-GFP), and thymidylate synthase (TMS-GFP) are excluded from the phage nucleus (Fig. 2E; [11]). At late stages of PhiPA3 infection, bouquets also weakly excluded fluorescence from each of these GFP fusion proteins, which was demonstrated with a line-plot of fluorescence intensity drawn through the long axis of the cell (Fig. 2E, green arrows). Although the extent of the exclusion from the bouquets was not as strong as exclusion from the nucleus, the ability to weakly exclude these proteins as well as soluble GFP and mCherry (Fig. 2F, Fig. S2) suggests that bouquets are densely packed with phage particles that prevent free diffusion of these cytoplasmic proteins through the structures.

To provide further support for this model and better understand how these structures form, we performed a series of co-localization experiments. First, we simultaneously examined the localization of tails (gp165-GFP) or capsids (gp136-GFP) with the phage nucleus shell protein (mCherry-gp53) (Fig. 3A & 3B). Capsids (gp136-GFP) assemble and dock on the nucleus at 45 mpi (Fig. 3B), then colocalize with DNA rings on either side of the nucleus by 70 mpi (Fig. 3B, see also Fig. 2A). This result of capsids co-localizing with phage DNA was also observed with 201Phi2-1 in the small percentage of infected cells where bouquets formed (Fig. S3). In comparison, the tail sheath protein gp165-GFP was uniformly distributed until 60 mpi (Fig. 2B), at which point it re-localized into foci inside DAPI stained rings on both sides of the nucleus (Fig. 3A). Next, we visualized the tail sheath protein (gp165-GFP) together with the capsids (gp136-mCherry) and found that tail sheath fluorescence occurred inside the ring of capsids (Fig. 3C, D). To study the relative timing of capsid and tail assembly, we simultaneously visualized capsid gp136-mCherry and tail sheath protein gp165-GFP during infection using time-lapse microscopy (Fig. 3E, Fig. S4). In the example shown in Figure 3E, capsids (red foci) leave the nucleus and migrate to form bouquets nearly simultaneously to the accumulation of tail proteins at the same sites flanking the phage nucleus. Most of the tail sheath protein gp165-GFP accumulates into two foci within approximately 4.5 minutes (270 s) (Fig. 3E).

**Figure 3.**
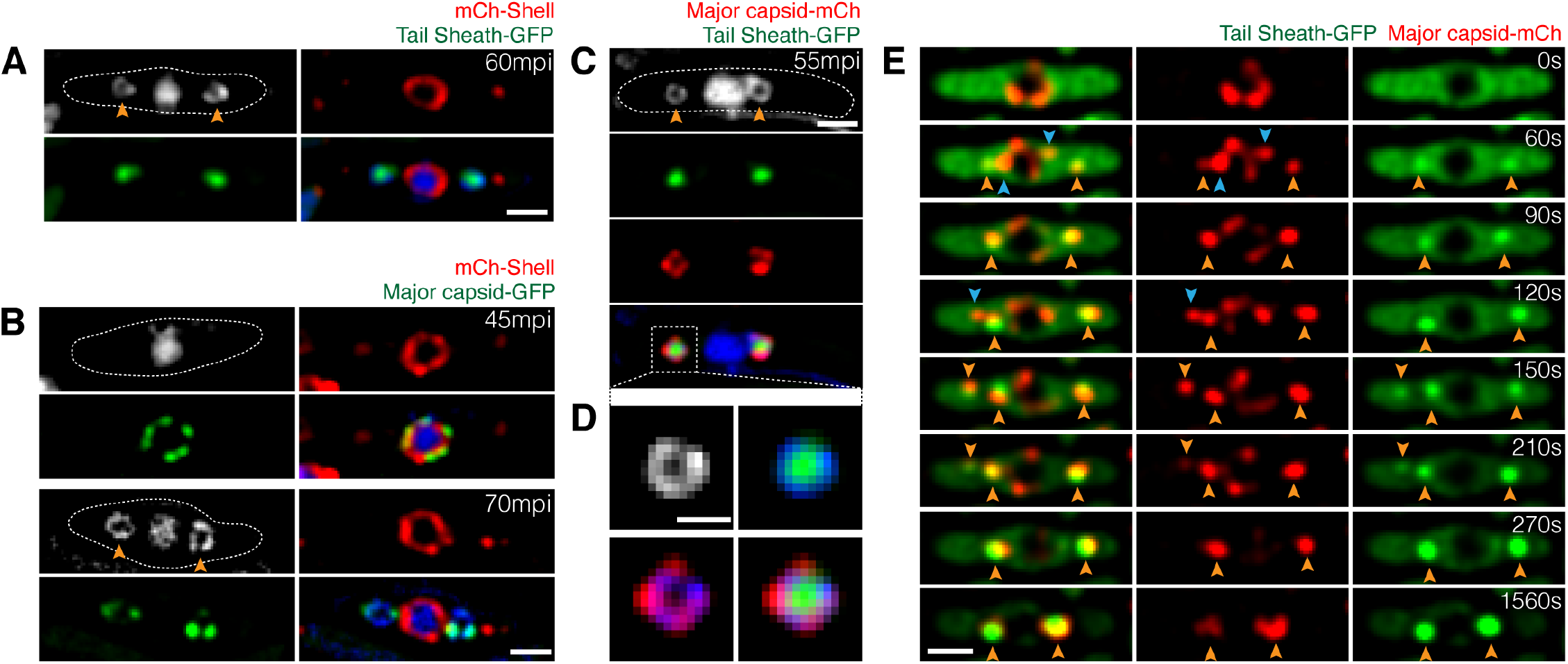
Temporal and spatial assembly of capsids and tails relative to the phage nucleus and bouquets. *P. aeruginosa* cells were grown on an agarose pad and the fusion protein was induced by arabinose at indicated concentration; 0.025% for mCherry-shell with tail sheath-GFP and mCherry-shell with major capsid-GFP, and 0.05% for major capsid-mCherry with tail sheath-GFP, prior to infection with phages PhiPA3. DNA was stained by DAPI (blue or gray). Dashed lines indicate the border of the cells. (A) Fluorescence images of cells expressing mCherry-tagged shell (gp053; red) and GFP-tagged tail sheath (gp165; green) infected with phage at 60 mpi. (B) Fluorescence images of cells expressing mCherry-tagged shell (gp053; red) and GFP-tagged major capsid (gp136; green) infected with phage at 45 and 70 mpi. (C, D) Fluorescence images of cells expressing mCherry-tagged major capsid (gp136; red) and GFP-tagged tail sheath (gp165; green) infected with phage PhiPA3 at 55 mpi. The bouquet structure is magnified (D) to reveal that capsids (red) overlayed on DNA and tails (green) were accumulated inside the structure. (E) Time-lapse imaging of mCherry-tagged major capsid (gp136; red) and GFP-tagged tail sheath (gp165; green) during an interval of 1,560 s starting at 45 mpi in the phage-infected cells. Capsids first packaged DNA on the surface of the phage nucleus and detached from the nucleus without tails (blue arrows). Mature capsids in cytoplasm later co-assembled with tails and formed bouquets (orange arrows). Scale bars: 1µm.

Taken together, our fluorescence microscopy experiments suggest that after DNA packaging, capsids and tails join and form bouquets late during the maturation process. To visualize this late step of the phage assembly pathway at higher resolution, we used cryo-electron tomography (cryo-ET). Cryo-ET can visualize samples that are thinner than ∼500 nm. In order to visualize PhiPA3-infected cells, which are over 1 um thick, we used cryo-focused ion beam (FIB) milling to generate ∼200 nm slices of infected cells. In samples collected at 70 mpi, we typically observed two clusters of capsids arranged in a spherical pattern (Fig. 4B, green). In the cell shown in Figure 4, one large cluster on the left of the phage nucleus contained eleven DNA-filled capsids (dark) arranged in a horseshoe pattern, in agreement with our fluorescence microscopy results (Fig. 1C). The distance from the edge of the capsid clusters to the edge of the phage nucleus (∼200 nm) also agreed with measurements from fluorescence microscopy. Of the eleven clustered capsids within the lamella, three appear to lack tails, but this could be due to FIB milling, which generated a 150 nm slice that may not have fully captured all capsids. Ribosomes (Fig. 4B, yellow) were excluded from the bouquets, in agreement with fluorescence microscopy images showing exclusion of ribosomal protein L28 (Fig. 2E). The opening of the horseshoe-shaped arrangement of capsids contained a dense material of unknown composition (Fig. 4B, grey). On the right side of the phage nucleus in Figure 4, four DNA filled capsids without tails are arranged in a tight grouping and, based upon its size, appeared to be a cluster at an early stage of formation. We also observed partially assembled tails unconnected from capsids and ranging in length from 85 to 140 nm (Fig. 4B, light blue).

**Figure 4.**
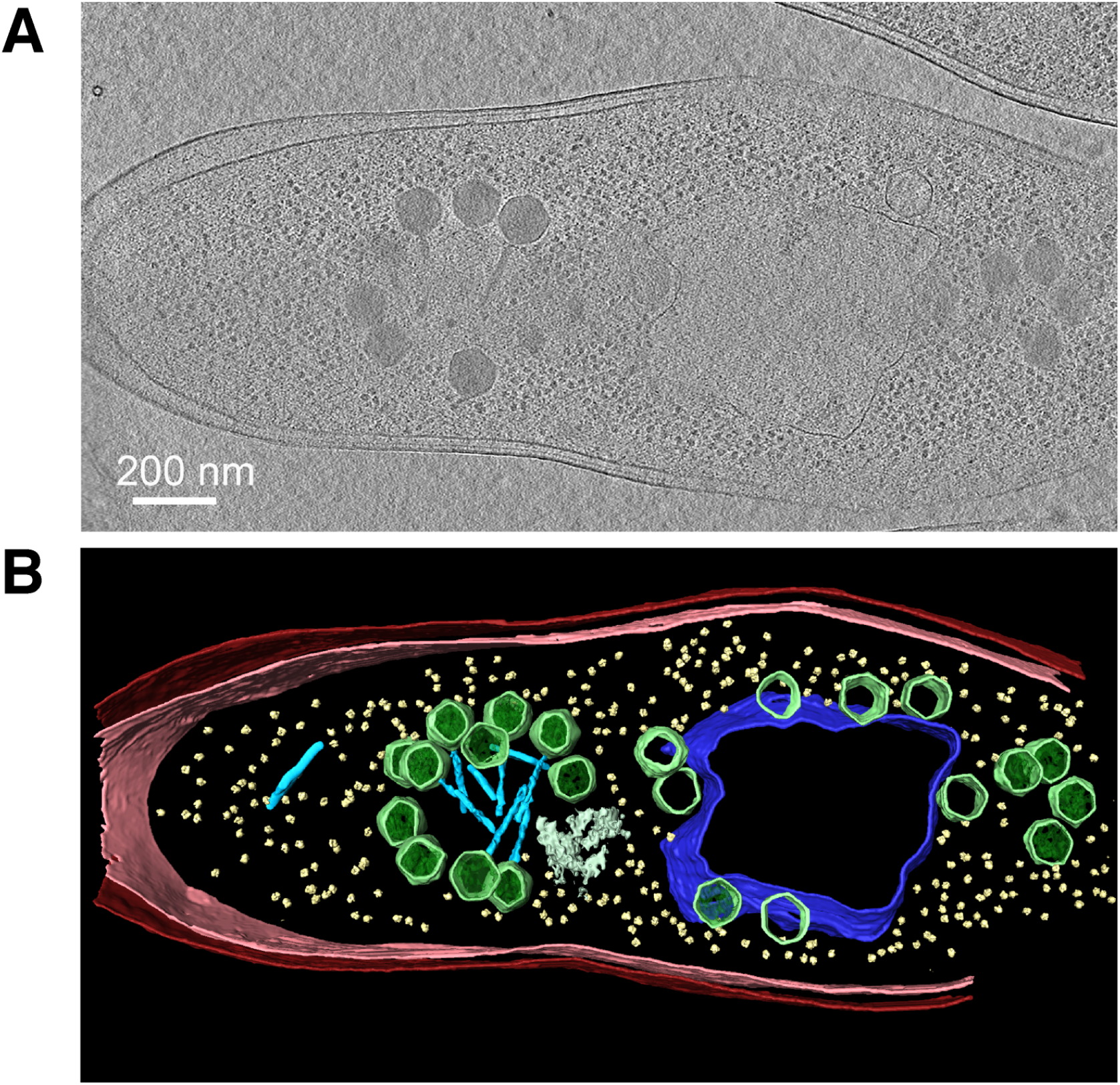
Cryo-electron tomography of phage PhiPA3-infected *P. aeruginosa* at 75 mpi reveals that the bouquet structure is made of mature phage particles with capsids facing outward and tails facing inward. (A) A slice through a tomogram of a cryo-focused ion beam–milled phage-infected cell at 75 mpi. (B) Segmentation of the tomogram in (A) showing extracted structures, including the shell (dark blue), capsids (green), tails (light blue), ribosomes (yellow), cytoplasmic membrane (pink), and outer membrane (red). The bouquet structure observed in cryo-electron tomography is horseshoe-shaped similar to that observed in 3D-SIM (Figure 1C). Scale bar: 200 nm.

Next, we attempted to identify determinants that affect bouquet formation or position relative to the phage nucleus. To determine if bouquet formation or positioning relies upon the PhuZ spindle, we studied PhiPA3-infected cells expressing PhuZD190A, a catalytically defective PhuZ mutant which behaves as a dominant negative. This mutant can bind GTP but is incapable of GTP hydrolysis. When expressed in PhiPA3-infected cells, monomers of PhuZD190A co-assemble with wildtype PhuZ produced by the phage to form long static filaments that are unable to properly position the phage nucleus at midcell or traffic capsids to it [11, 21]. Instead, the phage nucleus is mis-positioned, frequently appearing near the cell pole (Fig. 5A, bottom). Like the phage nucleus, the bouquets, which remain small puncta instead of rings as discussed below, are also mispositioned near the cell pole (Fig. 5A). However, they remain located at the same fixed distance of approximately 0.9 µm (n=109) from the center of the nucleus as in wildtype cells (0.9 µm, n=182) (Fig. 5G). These results suggest that bouquets rely upon positioning cues from the phage nucleus that are independent of the PhuZ spindle.

**Figure 5.**
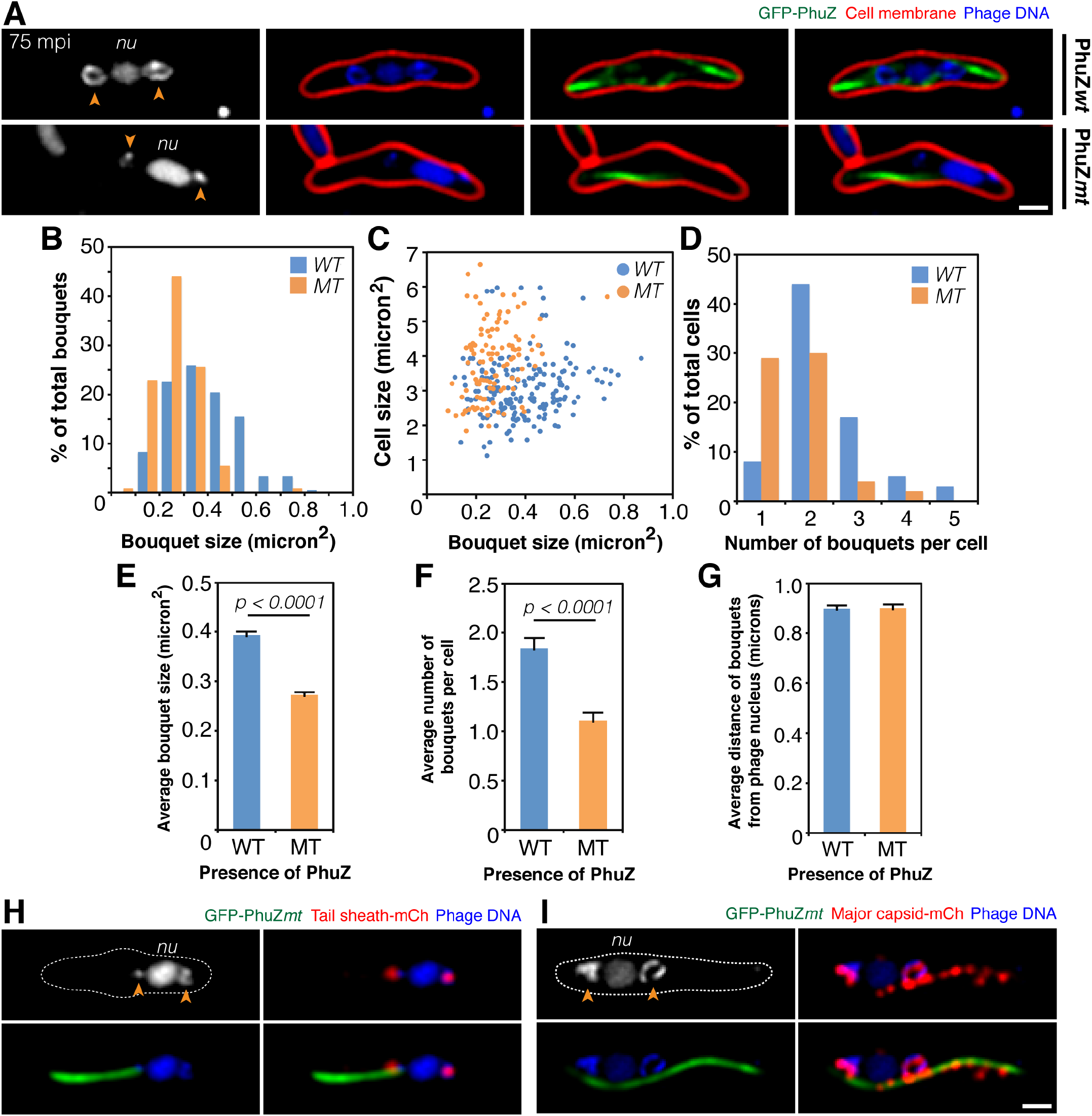
The formation of bouquets is impaired in the presence of mutant PhuZ filaments due to disrupted capsid trafficking. *P. aeruginosa* cells were grown on an agarose pad and the fusion protein was induced by arabinose concentration below the critical threshold for filament assembly; 0.025 - 0.1% for cells expressing wildtype and mutant PhuZ, prior to infection with phages PhiPA3. DNA was stained by DAPI (blue or gray). (A) Fluorescence images of infected cells expressing either wildtype GFP-tagged PhuZ (green; top panels) or catalytically defective GFP-tagged PhuZ (green; bottom panels) at 75 mpi. Cell membrane was stained red by FM4-64. Arrows indicate bouquets that locate adjacent to the nucleus (*nu*). The horseshoe-like structures were observed in the presence of wildtype PhuZ (top panel) but they appear smaller in the presence of mutant PhuZ (bottom panel). (B-G) Graphs for comparative analyses of bouquet structures between the cells expressing wildtype and mutant PhuZ; the percentage of bouquets at various sizes (B), bouquet size versus cell size (C), the percentage of cells containing different number of bouquets (D), average bouquet size (E), average number of bouquets per cell (F), average distance of bouquets from the phage nucleus (G). The average size and number of bouquets in the presence of mutant PhuZ are significantly lower (p < 0.0001) than those in the presence of wildtype PhuZ. Data were collected from the infected cells at 75 mpi from at least three different fields and are represented as mean ± SEM (n; all cells = 100, WT = 182 bouquets, and MT = 109 bouquets). (H-I) Fluorescence images of cells expressing catalytically defective GFP-tagged PhuZ (green) and either mCherry-tagged tail sheath (red; H) or mCherry-tagged major capsid (red; I) infected with phage PhiPA3 at 75 mpi. In the presence of mutant PhuZ, tail sheaths are able to localize to the bouquets, but capsids are trapped along the static filaments resulting in a defective bouquet structure. Scale bars: 1µm.

While bouquet positioning is independent of the PhuZ spindle, bouquet size and number decreased in the presence of the mutant. At 75 mpi, the majority (92%, n=100) of infected cells expressing wildtype PhuZ contained 2 or more bouquets while ∼8% contained a single bouquet (Fig. 5D). In contrast, in cells expressing PhuZD190A, ∼35% (n=100) of infected cells contained two or more bouquets and ∼30% contained a single bouquet at the same time point of infection (Fig. 5D), resulting in a small but significant decrease in the overall average number of bouquets per cell observed (Fig. 5F). The average bouquet size was also smaller, measuring 0.27 µm^2^ (n=109) in the cells expressing PhuZD190A compared to 0.39 µm^2^ (n=182) for the wildtype PhuZ-expressing strain (Fig. 5B, 5C, 5E). The decrease in bouquet size and number is likely due to the PhuZD190A mutant inhibiting capsid trafficking to the nucleus and therefore phage DNA packaging that precedes bouquet assembly [21]. To determine if this was the case, we localized capsids and tail proteins in the presence of the PhuZD190A-GFP mutant at late stages of infection. In wildtype cells, bouquets form on each side of the nucleus with tail proteins localized inside a sphere of capsid proteins (Fig. 2 & 3), with filaments of the wildtype PhuZ spindle extending past them to meet the phage nucleus at midcell (Fig. S5). In contrast, in the presence of the PhuZD190A-GFP mutant, tails were localized properly inside the bouquets (Fig. 5H), while many capsids were associated with the sides of the PhuZ filaments (Fig. 5I). These results suggest that, in the presence of the PhuZD190A-GFP mutant, bouquets decrease in size and number due to a failure of capsids to be transported to the phage nucleus, although bouquet positioning is largely independent of the PhuZ spindle. However, further work must be done to determine how the phage designate and maintain bouquet position.

Our discovery of a highly organized assemblage of phage particles inside bacteria late during infection demonstrates that nucleus-forming jumbo phages are capable of surprisingly complex subcellular organization throughout their entire replication cycle. Our updated model for jumbo phage replication, shown in Figure 6, begins with the attachment of phage to the host cell and injection of viral DNA. The expression of the nuclear shell protein occurs immediately after injection to form an enclosure around phage DNA that provides a safe compartment for DNA replication to occur. Bacterial chromosomal DNA is degraded, providing nucleotides for DNA synthesis and eliminating the ability of the cell to mount any further defense based on gene expression. Inside the phage nucleus, phage-encoded RNA polymerases transcribe viral mRNA that is exported by unknown mechanisms to the cytoplasm. Some of the proteins required for phage DNA replication, repair, recombination and transcription are selectively imported into the phage nucleus while phage structural proteins, metabolic enzymes and ribosomes remain in the cytoplasm. Viral capsid proteins assemble on the membrane midway through the infection cycle and are transported by treadmilling PhuZ filaments to the phage nucleus, where they dock to initiate DNA packaging. Treadmilling filaments rotate the phage nucleus to allow capsids to be evenly distributed on its surface. Once capsids are full of viral DNA, they co-assemble with tails and form bouquets containing what appear to be mature particles. Remarkably, these large assemblages of particles occur at a specific distance from the phage nucleus even when the nucleus is severely mispositioned due to expression of a catalytically dead PhuZ mutant. While our results suggest that the PhuZ spindle is likely not essential for their formation, we cannot completely rule out a role for the spindle in bouquet production since the PhuZ protein is also involved in delivering capsids to the nucleus.

**Figure 6.**
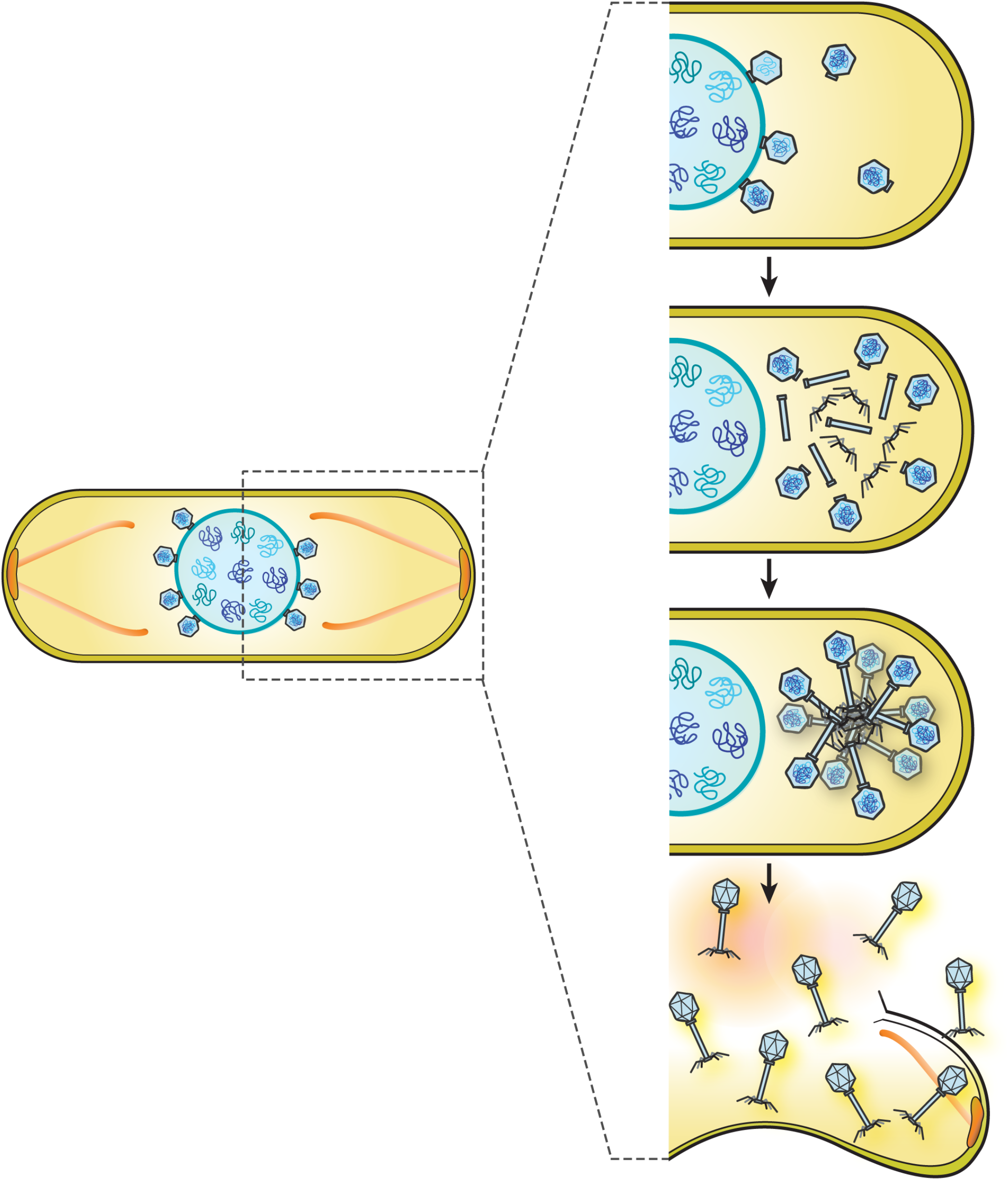
Model of bouquet assembly for maturation process of *Pseudomonas* jumbo phages. After procapsids have packaged DNA at the surface of the phage nucleus, mature capsids detach and localize in the cell cytoplasm. Tail components later assemble, then co-assemble with the DNA-filled capsids to form mature particles, which accumulate together with the capsids pointing outward while tails point inward to form a structure resembling a bouquet of flowers. The bouquet is able to grow in size and increase in number during late infection until the host cell lyses.

Assembly of viruses in eukaryotes typically occurs in defined subcellular compartments referred to as assembly factories or virus assembly centers (VACs) [1, 10]. For example, many eukaryotic viruses complete their final assembly at the plasma membrane, where high concentrations of viral proteins accumulate before being incorporated to form the mature virion. We now show that late during the lytic replication cycle, nucleus-forming bacteriophages also form large viral assemblages in defined subcellular compartments. Localized at fixed distances on either side of the phage nucleus, bouquets grow in size over time as new phage particles are added, demonstrating that their formation is spatially and temporally regulated. Bouquets exclude ribosomes and soluble enzymes such as GFP from their centers, suggesting that they represent a bona-fide subcellular compartment whose local physiology is distinct from the surrounding cytoplasm. As observed in eukaryotic virus assembly centers, phage bouquets might increase the efficiency of viral particle production, although this hypothesis remains to be tested.

We do not currently understand how bouquets form but propose two possible hypotheses. In one scenario, capsids are filled with DNA, leave the nucleus surface, and attach to tails that have assembled in the cytoplasm. The mature particles then accumulate in bouquets. In one variation of this model, clusters of tails assemble adjacent to the nucleus, followed by the attachment of DNA filled capsids. Alternatively, tail-less capsids might assemble in a spherical configuration adjacent to the nucleus, followed by assembly and attachment of tails in the bouquets. We cannot currently distinguish between these models. Although the role of bouquet formation in phage replication is unclear, it is tempting to speculate that they are viral assembly centers that increase the efficiency of phage particle maturation. As they exclude examined host cell proteins (Fig. 2E), the bouquet structure may also protect the phage particles from host proteases or other defenses that might damage tail fibers and reduce their infectivity. However, we note that bouquets are not observed in every cell during PhiPA3 infection and are less frequently observed during infections by related phages PhiKZ and 201Phi2-1. Therefore, it seems likely that these assemblages are not essential for these phages to reproduce. In relatively rare cases, previous electron microscopy studies examining phage lysates have noticed clusters of phage in a bouquet structure outside of cells, but it was unclear if these structures were formed inside of cells as part of phage production or if they resulted from attachment to a common substrate after cell lysis [37-39]. Here we show that phage bouquets form naturally *in vivo* during replication of *Pseudomonas* jumbo phages. Taken together, our results expand our knowledge of viral subcellular organization and suggest that viral assembly and maturation is spatially organized in viruses across kingdoms.

## Methodology

### Strains, growth condition and phage preparation

*P. chlororaphis* strain 200-B, *P. aeruginosa* strain PA01 and PA01-K2733 (Pump-knockout) were cultured on solid Hard Agar (HA) (Serwer et al., 2004) and Luria-Bertani (LB) media, respectively, and incubated at 30°C overnight. High-titer phage lysates of each phage: 201Phi2-1, PhiPA3, and PhiKZ, were prepared by infecting its corresponding bacterial host culture with 10 µl of phage lysate and then incubating for 15 minutes at room temperature. 5 mL of HA (0.35%; phage 201Phi2-1) or LB top agar (0.35%; phage PhiPA3 and PhiKZ) was mixed with the phage-infected cultures and was poured over a HA or LB plate. After the agar was solidified, the plates were then incubated upside-down at 30°C overnight. The following day, the plates that formed nearly confluent lysis (web lysis) were flooded with 5 mL of phage buffer and allowed to sit at room temperature for 5 hours. The phage lysates were then collected, clarified by centrifugation at 15,000 rpm for 10 minutes, and stored at 4°C with 0.01% chloroform.

### Plasmid constructions and bacterial transformation

Phage genes of interest were first amplified from high-titer phage lysates using PCR amplification. Each amplicon was then ligated into the linearized backbone pHERD-30T to generate a recombinant plasmid via isothermal assembly using NEBuilder^®^ HiFi DNA Assembly Cloning Kit (Cat. No. E5520S). The recombinant plasmid construct, as listed in Table 1, was later transformed into E. coli DH5a in which the transformants were plated on LB supplemented with gentamycin sulfate (15 µg/mL). Constructs were confirmed by DNA sequencing and subsequently introduced into indicated organisms by electroporation, resulting in strains listed in Table S1. The transformants were selected on LB supplemented with gentamycin sulfate (15 µg/mL).

**Table 1.**
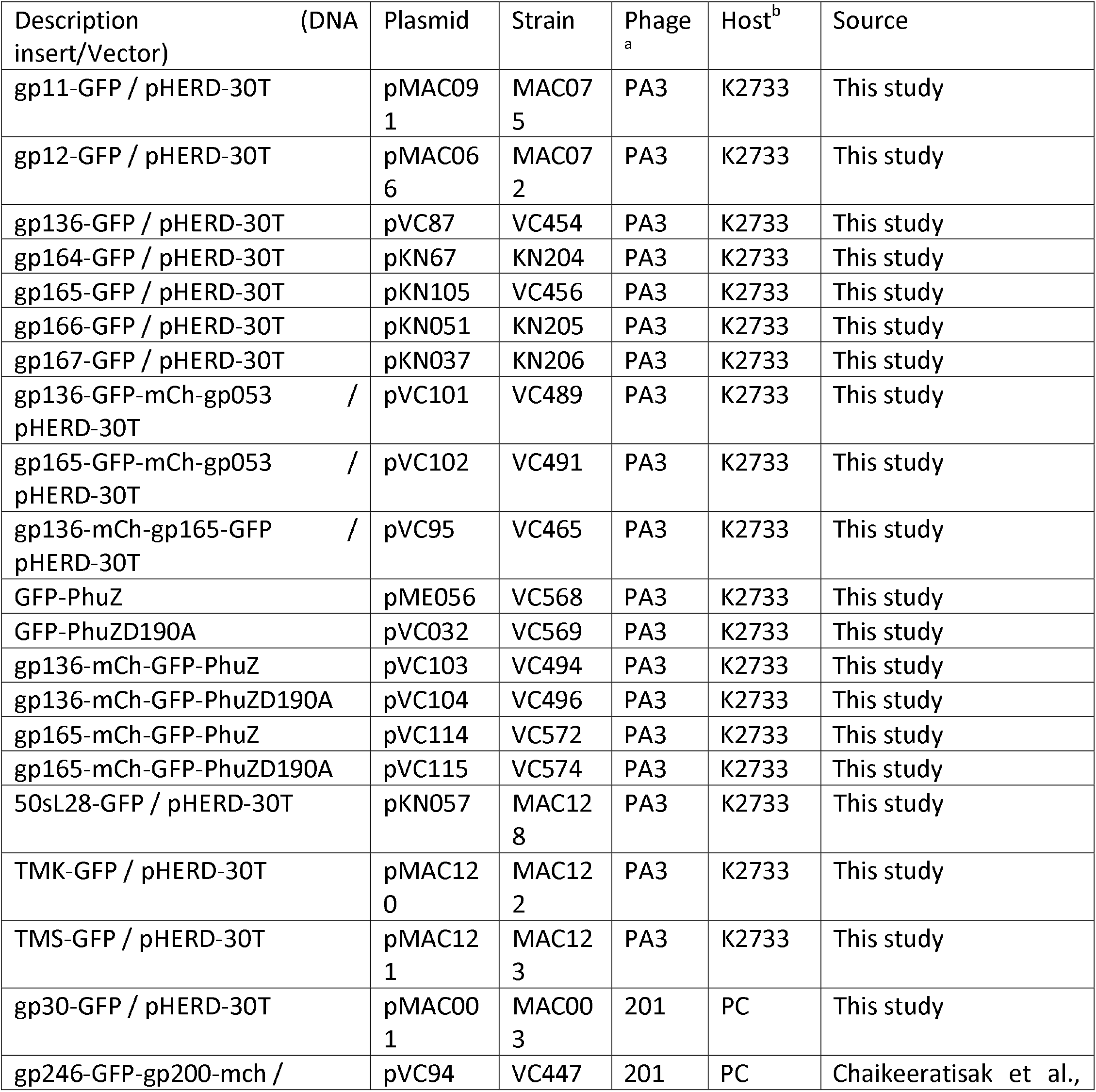
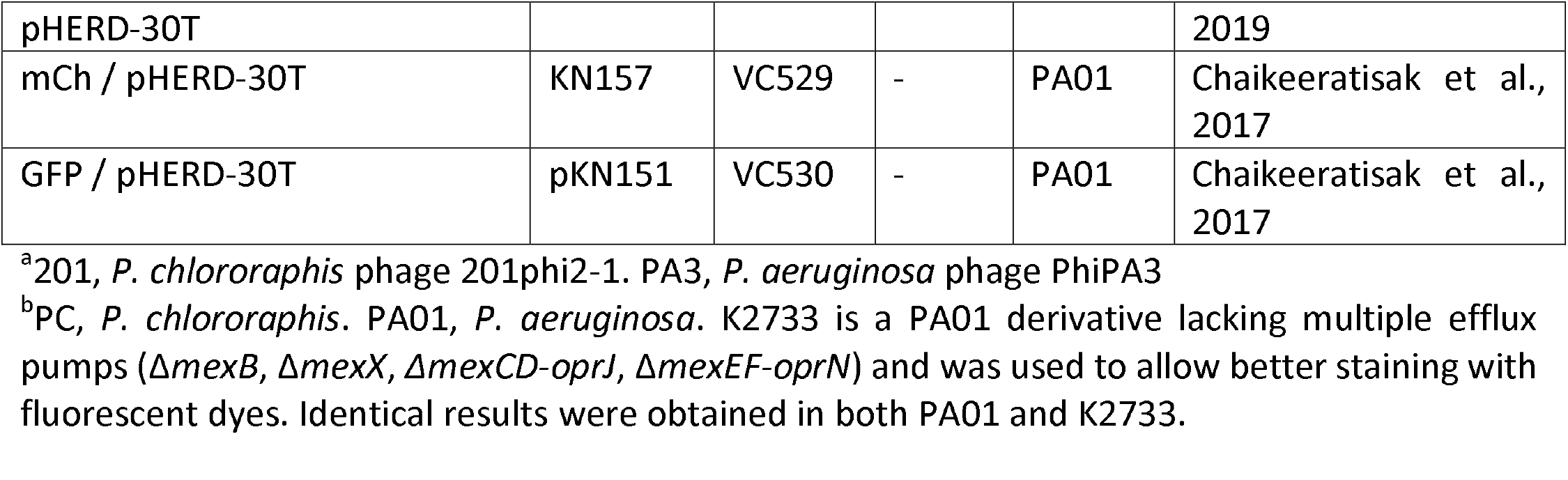
Plasmids and strains used in this study

### Fluorescence microscopy and single cell-infection assay

Bacterial cells were inoculated on 1.2% agarose pads prepared on concavity slides. Each pad was supplemented with arabinose concentrations as indicated to induce the protein expression, 1 µg/mL FM4-64 for cell membrane staining, and 1 µg/mL DAPI for nucleoid staining (Pogliano et al., 1999). *P. chlororaphis* strain 200-B and *P. aeruginosa* strain PA01-K2733 were grown on pads in a humid chamber at 30 °C for 3 hr and at 37 °C for 2 hr, respectively. To initiate the phage infection, 5 µl of high-titer phage lysate (10^8^ pfu/mL) was added onto the cells on agarose pads, and the pad was then incubated at 30°C (201Phi2-1) or 37°C (PhiPA3 and PhiKZ) to allow the infection to proceed. At desired time point, a coverslip was put on the slide and fluorescence microscopy was then performed.

The DeltaVision Spectris Deconvolution Microscope (Applied Precision, Issaquah, WA, USA) was used to visualize the cells. For static images, the cells were imaged for at least 8 stacks in the Z-axis from the middle focal plane with 0.15 µm increments. For time-lapse imaging, the cells were imaged from a single stack at the focal plane at selected intervals as indicated using ultimate focusing mode. Collected images were further processed by the deconvolution algorithm in theDeltaVision SoftWoRx Image Analysis Program and later analyzed in Fiji (Schindelin et al., 2012).

### 3D-SIM super-resolution microscopy

*Pseudomonas* cells were grown and infected with corresponding phages as described above for live-cell conditions. At desired 90 mpi, the infected cells were fixed with paraformaldehyde and glutaraldehyde, then washed by 1X PBS as previously described in Chaikeeratisak et al., 2017. The fixed cells were incubated with 1 µg/mL DAPI to stain all DNA. Applied Precision/GE OMX V2.2 Microscope was then used to image and collect raw data. Raw data were taken by SI-super-resolution light path to collect 3 µm thickness of samples with 125 nm increments in the z-axis with compatible immersion oils (Applied Precision). 3D-structured illumination microscopy (SIM) images were then rendered by standard OMX SI reconstruction parameters in DeltaVision SoftWoRx Image Analysis Program, and later analyzed in Fiji (Schindelin et al., 2012).

### Quantification and data analysis

All experiments for quantification and data analysis were done from at least 3 independent biological experiments. All data are shown as mean values or mean ±SEM, as indicated in figure captions. The number of cells or phage bouquets and size/distance measurement were manually counted in Fiji (Schindelin et al., 2012). Pairwise comparison between the data was conducted for unpaired data with unequal variance using Student’s t test. A *p* value less than 0.05 indicates a significant difference. All statistical analysis, data processing, and data presentation were conducted using KaleidaGraph and Microsoft Excel.

### Tomography sample preparation and Cryo-FIB electron microscopy data acquisition

*P. aeruginosa* cells were infected with ΦPA3 as outlined in sample preparation and at 70 mpi, cells were scraped off the agarose pad and suspended in ¼ LB media. Holey carbon coated QUANTIFOIL® R 2/1 copper grids were glow discharged using Pelco easiGlowTM glow discharge cleaning system. Manual plunging and cryo-FIB milling using Scios dual beam (Thermo Fisher Scientific) were performed as detailed in [11, 21]. Tilt-series were collected from typically −60° to +60° with 2° tilt increments using SerialEM in a 300-keV Tecnai G2 Polara microscope (Thermo Fisher Scientific) equipped with post-column Quantum Energy Filter (Gatan) and a K2 Summit 4k x 4k direct detector camera (Gatan) [40]. Images were recorded at a nominal magnification of 27,500 (pixel size, 0.748 nm) with the dose rate of 10-12 e/physical pixel at the camera level. The total dose in the tomogram in Fig 4A was ∼51 e^-^/A^2^ with −5 μm defocus. A total of 10 tomograms of *P. aeruginosa* infected with ΦPA3 at 70 mpi were collected from 5 cryo-FIB milled lamellae with bouquet-like structures visible in 4 of these. Tilt-series were reconstructed in IMOD using patch tracking method [41]. Membranes and capsid shells were semi-automatically segmented with TomoSegMemTV that were then manually refined with Amira software (Thermo Fisher Scientific) [42]. EMAN2 was used to manually pick the ribosomes, which were then subsequently averaged, classified and placed back in the tomogram using Dynamo [43, 44].

## Acknowledgements

This research was supported by National Institutes of Health grants GM104556 (J.P.) and GM129245 (J.P. and E.V.), and a National Science Foundation MRI grant DBI 1920374(E.V). We acknowledge the use of the UC San Diego cryo-EM facility, which was built and equipped with funds from UC San Diego and an initial gift from Agouron Institute, and the San Diego Nanotechnology Infrastructure (SDNI) of UCSD, a member of the National Nanotechnology Coordinated Infrastructure, which is supported by the National Science Foundation (Grant ECCS-1542148).

## Notes

### Competing Interest Statement

The authors have declared no competing interest.

