## Supplementary Material for "Subcellular Organization of Viral Particles During Maturation of Nucleus-Forming Jumbo Phage"

#### Supplemental Figure Legends

**Figure S1.** Fluorescent micrographs of uninfected control cells. The cells were grown on agarose pads, incubated at 30°C for 3 hours, and were induced by indicated arabinose concentration to express fluorescent protein fusions. Membranes were stained with FM 4-64 (red) and DNA with DAPI (blue). Cover slips were put on immediately before the microcopy and images were then collected. PA3-gp12, PA3-gp136, PA3-gp164, PA3-gp165, PA3-gp166, PA3-gp167 appeared uniformly distributed throughout the cells in the absence of infection whereas PA3-gp11 formed small foci. Scale bar, 1 micron.

**Figure S2**. Fluorescent micrographs of PA3 infected *P. aeruginosa* cells expressing fluorescent protein; GFP and mCherry. The cells were grown on agarose pads and were induced by 0.2% arabinose to express the fluorescent protein prior to infection. At late time point, phage bouquets were assembled and partially excluded fluorescent proteins suggesting that the physiology inside phage bouquet is distinct from cell cytoplasm. Scale bar, 1 micron.

**Figure S3.** Fluorescent micrographs of 201Phi2-1 infected *P. chlororaphis* cells expressing major capsid (gp200)-mCherry and internal head protein (gp246)-GFP showing phage maturation of phage 201Phi2-1 *in vivo*. (A, B). At late time point after DNA packaging, extranuclear staining structures we term “Phage Bouquet” (orange arrows) appear adjacent to the phage nucleus (nu). Major capsid (gp200, red) and the internal head protein (gp246, green) colocalize on the structure suggesting that the structures are comprised of the DNA packaged-mature capsids. Scale bar, 1 micron. The region inside the dashed box in (A) is magnified in (B) to more clearly show the colocalization of major capsid and the internal head protein on the phage bouquet. DAPI staining is shown in white in the top row panel A, in blue in the bottom row of panel A, and in white and blue in panel B. Scale bar, 0.5 micron.

**Figure S4.** Time-series of fluorescent micrographs of PA3 infected *P. aeruginosa* cells expressing major capsid (gp136)-mCherry and tail sheath (gp165)-GFP. The cells were grown on agarose pads and were induced by 0.05% arabinose to express fluorescent proteins. High titer phage PA3 lysates were added on top of the pad to initiate infection. Time-series throughout infection were imaged and collected. By 45 mpi, capsids assemble, package DNA at the phage nucleus, detach and localize in the cytoplasm. At this time point, capsids (red foci) did not appear to contain assembled tails (pink arrows). After 45 mpi, more capsids were incorporated into phage bouquets where tails localized inside and were surrounded by capsids (orange arrows). Scale bar, 1 micron.

**Figure S5.** Fluorescent micrographs of PA3 infected *P. aeruginosa* cells expressing wildtype GFP-PhuZ with either tail sheath (gp165)-mCherry (A) or major capsid (gp136)-mCherry (B) at late infection. The cells were grown on agarose pads and were induced by 0.025% arabinose to express the fluorescent proteins. High titer phage PA3 lysates were added on top of the pad to initiate infection. In the presence wildtype PhuZ filament, phage bouquets were located adjacent to the phage nucleus. Major capsids and tail sheath were localized and associated with the phage bouquets. Scale bar, 1 micron.

### Figure S1

#### Uninfected cell control
(induced with indicated arabinose concentration)

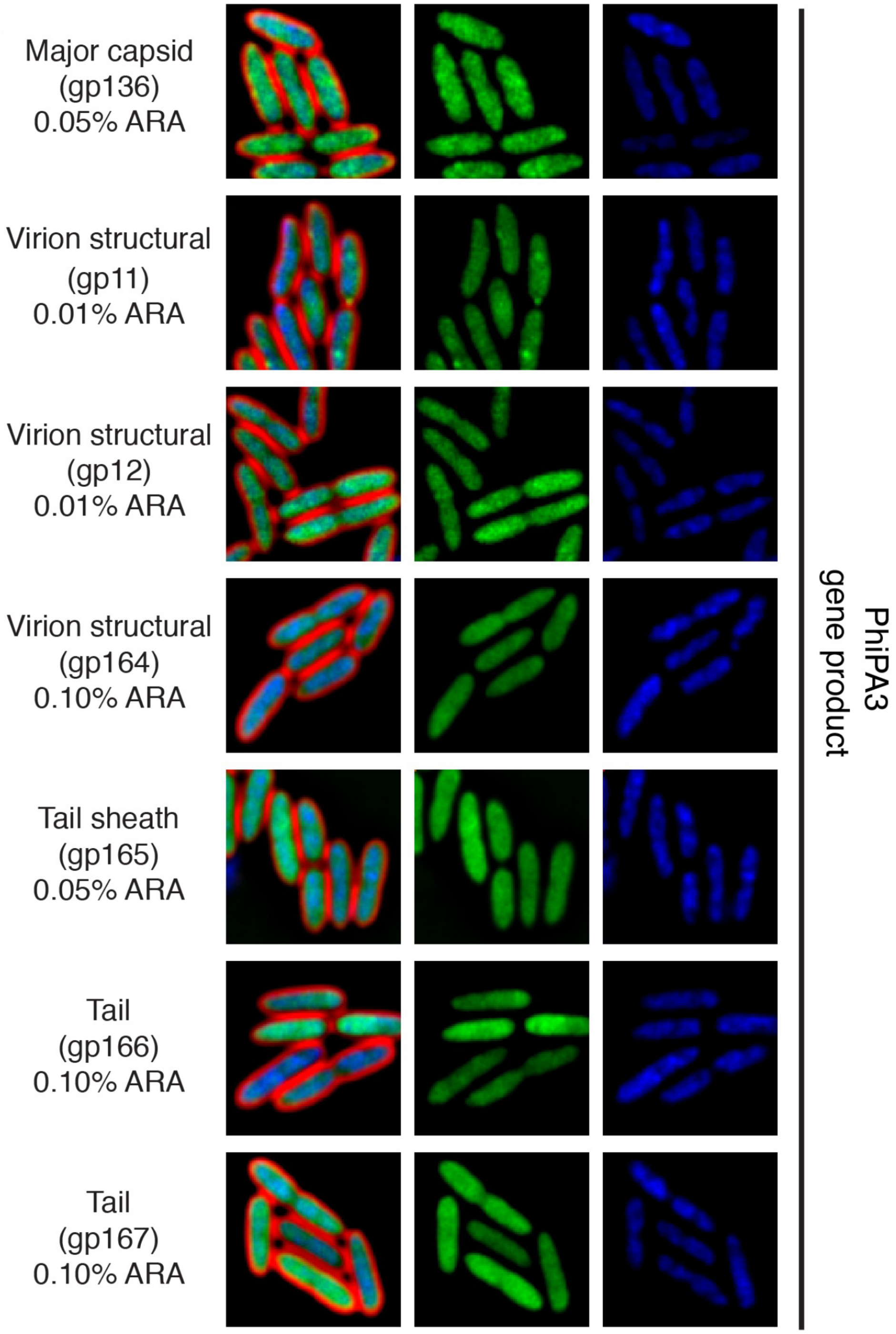

### Figure S2

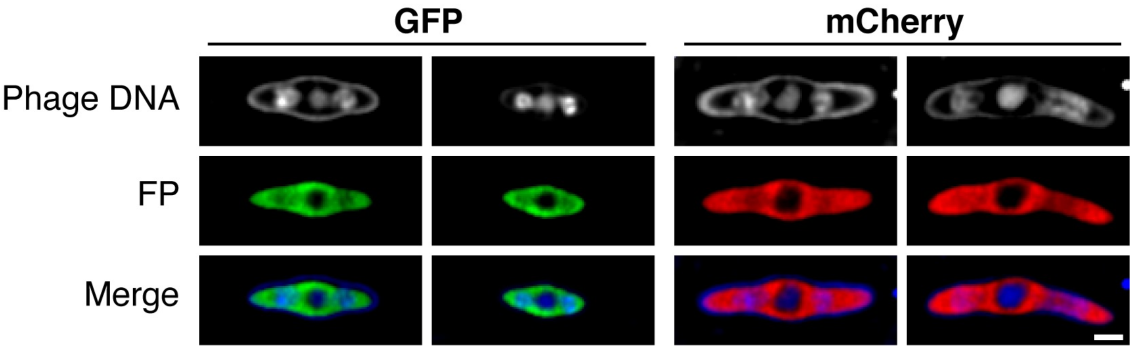

#### Figure S3

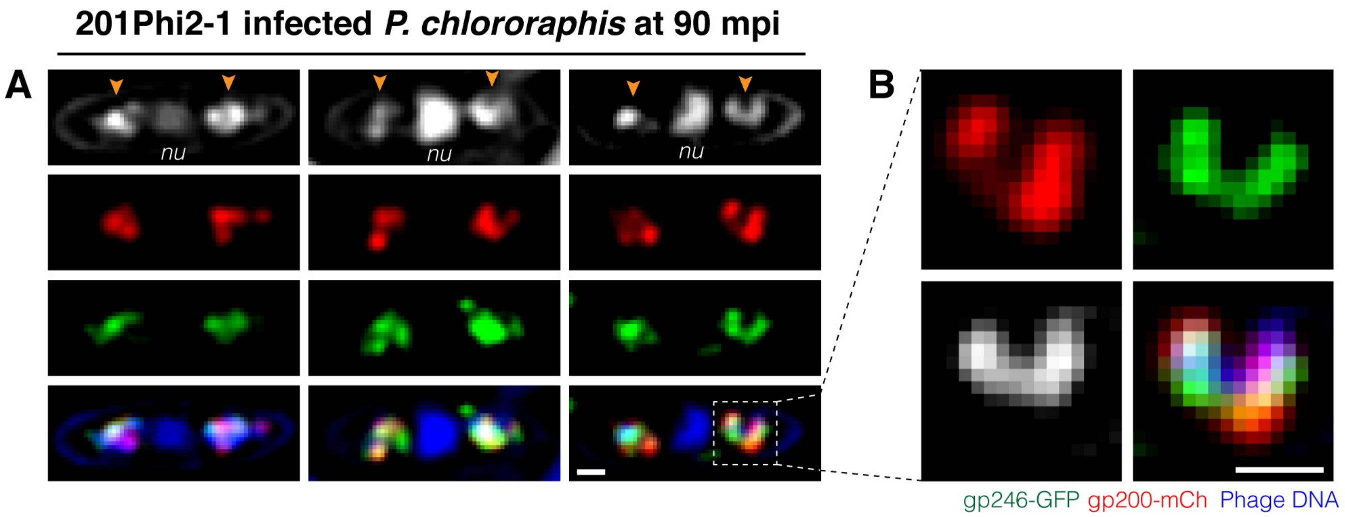
201Phi2-1 infected *P. chlororaphis* at 90 mpi
A
nu
nu
nu
B
gp246-GFP gp200-mCh Phage DNA

### Figure S4

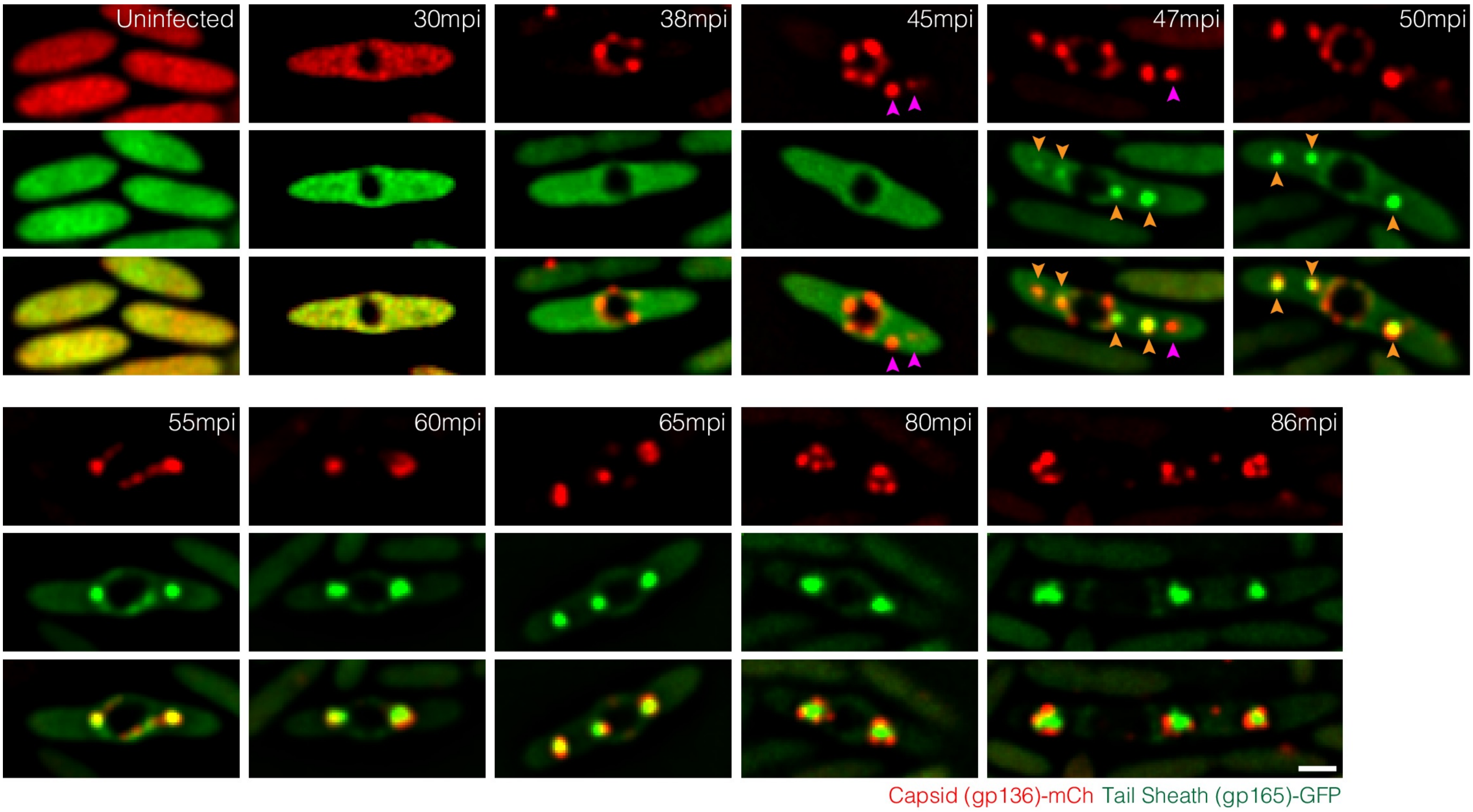

Uninfected
30mpi
38mpi
45mpi
47mpi
50mpi
55mpi
60mpi
65mpi
80mpi
86mpi
Capsid (gp136)-mCh Tail Sheath (gp165)-GFP

### Figure S5

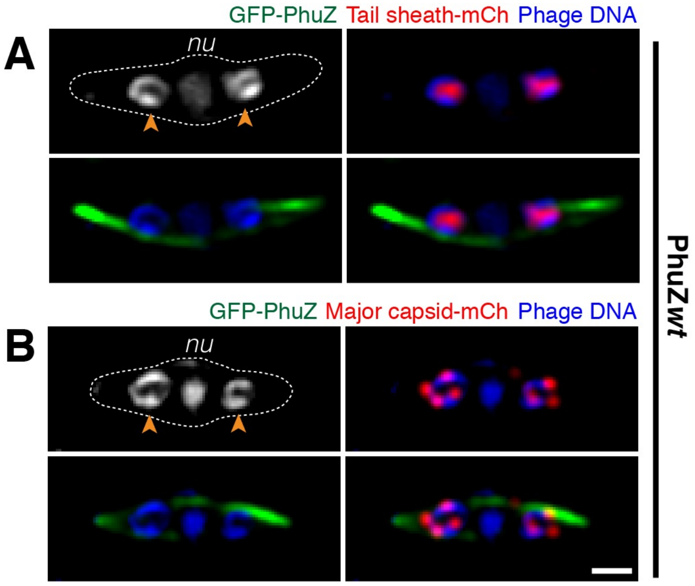

GFP-PhuZ Tail sheath-mCh Phage DNA
A
nu
GFP-PhuZ Major capsid-mCh Phage DNA
B
nu
PhuZwt
